# The Recombination Triplet State in the Far-Red Light Adapted Photosystem II is Located at the Chl_D1_ Site and Resides on the Red-Most Chlorophyll of the Reaction Center

**DOI:** 10.1101/2025.10.15.682561

**Authors:** Andrea Calcinoni, Anna Paola Casazza, Stefania Viola, Alessandro Agostini, Donatella Carbonera, Stefano Santabarbara

**Author notes:** To whom correspondence should be addressed: S.S.; D.C. These authors contributed equally (Andrea Calcinoni, Anna Paola Casazza).

## Abstract

The energetic limits of Photosystem II (PSII) photochemical reactivity required reconsideration after the discovery of far-red light acclimation responses in cyanobacteria. Insights on PSII functionality following the inclusion of the red shifted Chlorophylls *d* and *f* can be obtained by extending the current knowledge on spectroscopic and structural properties of its reaction center (RC). The photo-induced triplet states, that represent selective endogenous probes, were therefore investigated in far-red adapted PSII by magnetic resonance techniques. Zero-field splitting tensor analysis combined with spin-polarization dynamics arising from radical pair recombination unambiguously identify an intrinsically low-energy-absorbing chlorophyll participating to charge separation reactions. The triplet-*minus*-singlet (T–S) spectrum associated to the recombination triplet state, obtained by microwave selection, showed a sharp 725 nm bleaching demonstrating the dominant involvement of this red-shifted chlorophyll in the lowest RC exciton. Moreover, spectral simulations provided strong evidence in favor of its localization at the Chl_D1_ position, making it the most likely site of primary photochemistry.

---

Photosystem II (PSII) is a multi-subunit cofactor-binding complex that represents a key component of oxygenic photosynthesis, catalyzing the light dependent water splitting. The so-called core complex of PSII comprises four main chromophore-cofactor binding subunits: CP43 and CP47 that serve as a proximal light harvesting antenna to the photocatalytic reaction center (RC), whose cofactors are primarily coordinated by the D1/D2 heterodimer. In the vast majority of oxygenic phototrophs, Chlorophyll (Chl) *a* is the dominant pigment bound to the core complex, being active both in light harvesting and in photochemistry within the RC, in conjunction with Pheophytin (Pheo) *a* (Figure 1A). The nearly ubiquitous presence of Chl *a* in the RC of PSII has been linked to minimal energy requirements necessary for water splitting, in particular to the energy of its lowest singlet excited state that lays at approximately ∼1.8 eV with respect to the singlet ground state. This general view has become under scrutiny following the discovery of oxygenic phototrophs that contain lower energy absorbing Chls, such as Chl *d*, that replaces almost entirely and constitutively Chl *a* in *Acaryochloris marina*, and Chl *f*, that can replace about 10% of Chl *a* in cyanobacteria capable of the conditional Far-Red Light Photoacclimation (FaRLiP)^**1-2**^. The FaRLiP response entails a significant reshaping of the photosynthetic apparatus, leading to the replacement of some key subunits of PSI and PSII with specific isoforms^**3**^. It has been suggested that, in the far-red (FR) adapted PSII, together with the substitution of some Chl *a* molecules with Chl *f* in CP43 and CP47 to tune the light harvesting bandwidth, at least one of the Chl *a* participating in photochemical reactions within the RC is replaced by either Chl *f* or Chl *d*^**4**^. Investigations by steady state and time resolved optical spectroscopy^**5-8**^ also support the involvement of Chl *d/f* in PSII photochemical charge separation, with early modelling proposing them to be located either at the P_D1/D2_ dimer or at the so-called Chl_D1/D2_ sites^**5**^. The assignment of Chl_D1_ being Chl *d* has been favored by Cryo-EM structural studies^**9**^ and recent QM/MM investigations^**10**,**11**^. Although the assignments are also supported by indirect evidences, such as the identification of possible isoform-specific residues coordinating peripheral substituents of the Chl *d/f* chlorin macrocycles, the unequivocal identification of Chl species remains challenging when the structural coordinates are far from atomic resolution. Hence, the location and molecular identity of the Chl *d/f* molecules in the FR-PSII RC, and, consequently, the exact sites of photochemical and charge stabilization events, remain to be proven experimentally. Useful insights to address these issues can be obtained from the analysis of photo-induced triplet states. These species, which are sensitive and selective internal probes of the RC chromophores^**12-14**^, were here investigated by both Time-Resolved Electron Paramagnetic Resonance (TR-EPR) and Optically Detected Magnetic Resonance (ODMR) in the FR-PSII. Under reducing conditions, a triplet state displaying the characteristic electron spin polarization (esp) resulting from the radical-pair recombination mechanism, which is illustrated in the reaction scheme in Figure 1, was detected. The zero-field splitting (ZFS) parameters associated with this triplet state unambiguously demonstrate its localization on either a Chl *d* or a Chl *f* rather than Chl *a* cofactor. Moreover, this chromophore dominates the lowest energy state of the FR-adapted PSII RC, as evidenced by the maximal ground state bleaching at 725 nm of the microwave-induced triplet-*minus*-singlet (T–S) spectrum, red-shifted more than 40 nm (0.11 eV lower energy) relative to the canonical Chl *a*-binding PSII (Figure 1B). Spectral simulations assign this low-energy pigment to the Chl_D1_ site. Since the observed lowest energy triplet esp stems from a singlet state radical-pair precursor, it can be concluded that Chl *d/f* at Chl_D1_ is directly involved in primary photochemistry.

**Figure 1.**
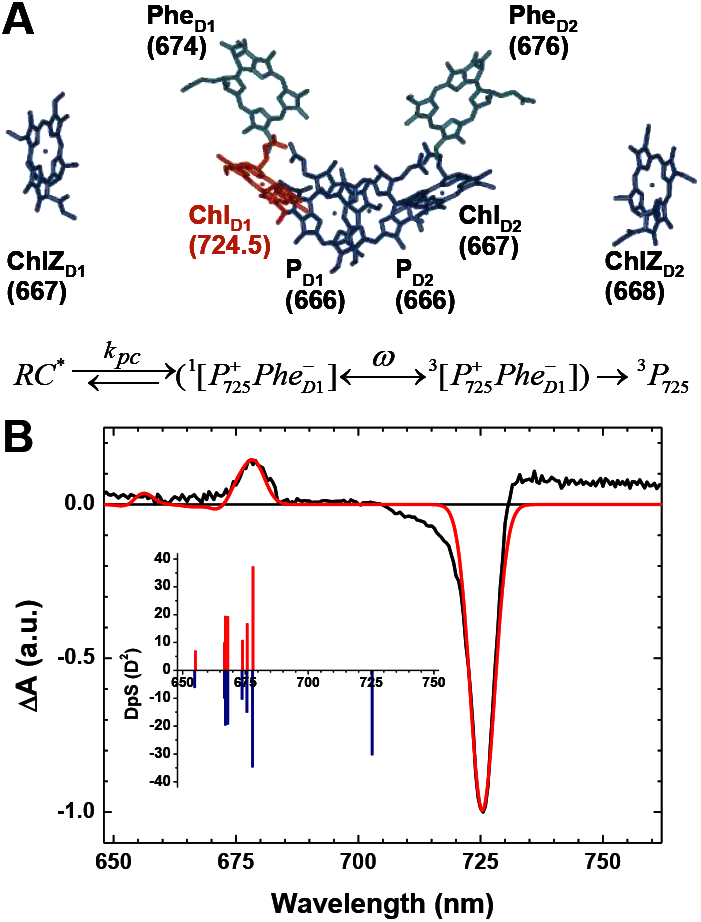
**A:** Chromophore arrangement in the FR-PSII RC (PDB 8EQM) and site energies (in nm) retrieved from T–S spectral simulations considering the photo-induced excited triplet state being populated by the recombination mechanism, as shown in the reaction scheme (where, *k*_pc_, photochemical charge separation rate; ω, singlet-triplet-mixing rate) and localized on the Chl_D1_. **B:** Comparison of the experimental T–S spectrum (black line) acquired upon microwave selection at 570 MHz (marked with a black arrow in Figure 2A) and the simulated one (red line), each normalized to the maximal bleaching. Inset: stick spectra displaying the eigenstates and associated populations for the system being either in the singlet (blue) or triplet (red) state. Further details on the spectral simulation are reported in SI.

Measurements were performed on the PSII core complex isolated from thylakoid membranes of *Chroococcidiopsis thermalis* PCC7203 grown under far-red illumination (750 nm, see SI for further detail). The FR-PSII sample was incubated with Na_2_S_2_O_4_ (10 mM) under anaerobic conditions and illuminated at room temperature, in order to fully reduce the PSII quinone acceptors, a condition that, in canonical PSII RC, promotes the recombination of charge-separated states to the triplet state. ODMR analysis on photo-induced FR-PSII triplets was initially conducted by Absorption Detected Magnetic Resonance (ADMR), taking advantage of the differences in the absorption maxima of Chl *a* and Chl *d/f*.

Figure 2A/B show the ^3^Chl *d/f* ADMR spectra detected at 725 nm, displaying maxima at 566 MHz in the │D│–│E│ transition and at 875 MHz, comprising a distinct shoulder at 890 MHz, in the │D│+│E│ transition. The marked asymmetry of the │D│+│E│ transition suggests the presence of two subpopulations. Gaussian deconvolution of both transitions indicates that the two triplet states have the following ZFS: │D│= 0.0239 cm^-1^, │E│= 0.0052 cm^-1^ and │D│= 0.0244 cm^-1^, │E│= 0.0053 cm^-1^. These correspond to a decrease in │D│ and parallel increase in the │E│ values with respect to the ZFS previously determined for ^3^Chl *d* either *in vitro*^**15**^ or bound to the PSI supercomplex of *A. marina*^**16**,**17**^. At the same time, the ZFS are not largely different from those attributed to a ^3^Chl *f* observed in the recombinant Chlorophyll *f* synthase (rChlF) enzyme (│D│= 0.0251 cm^-1^, │E│= 0.0051 cm^-1^)^**18**^. Nevertheless, the differences just discussed appear to fall within the ZFS tuning range that protein coordination can exert, as already demonstrated for the thoroughly investigated ^3^Chl *a*^**12**,**13**^. Then, at present, it is not possible to unambiguously assign the observed triplet to either Chl *d* or Chl *f*, whereas the involvement of ^3^Chl *a* can be excluded.

**Figure 2.**
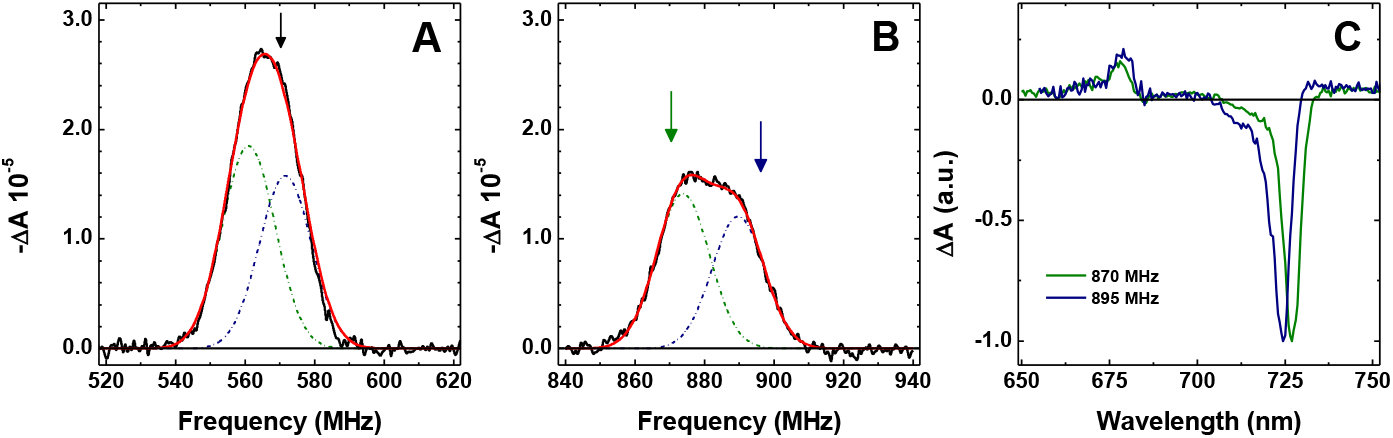
**A** and **B:** ADMR spectra recorded at 725 nm in the ^3^Chl *d/f* │D│–│E│ and │D│+│E│ transitions respectively (black lines), together with their decomposition (red lines) by a linear combination of two Gaussian sub-bands (dashed dotted green and blue lines). **C:** T–S spectra recorded upon microwave selection at 870 MHz (green line) and 895 MHz (blue line), preferential for the triplet populations indicated by the arrows in **B**. The spectra are normalized to their maximal bleaching. Experimental conditions: T= 1.8 K; Modulation Frequency= 33 Hz; mw power=0.5 W.

Consistently, the T–S spectrum obtained upon microwave selection at 570 MHz (│D│–│E│) displays a sharp ground state bleaching at 725.5 nm, accompanied by a relatively small amplitude positive absorption feature in the 670 – 685 nm window (Figure 1B). Figure 2C also reports the T–S spectra recorded for preferential selection of the two Chl *d/f* triplet populations discernible in the │D│+│E│ transitions, with pump-frequencies at 870 and 895 MHz. These T-S spectra show an overall similar shape to the one obtained at 570 MHz, but the exact position of the main bleaching shifts to 727 nm (for 870 MHz) and 724.5 nm (for 895 MHz), most likely indicating the presence of site-specific conformational, rather than chromophore occupancy, heterogeneity. It is worth noticing that the ^3^Chl *d/f* recorded in FR-PSII can also be observed in thylakoids pre-illuminated under reducing conditions (Figure S4), where remarkably similar ADMR and T–S spectra are observed, thereby excluding purification artefacts.

ADMR detection at 682 nm (Figure 3A/B) shows the presence in the system also of ^3^Chl *a* with maxima at 730.5 MHz and 994 MHz, corresponding to ZFS of │D│= 0.0288 cm^-1^, │E│= 0.0044 cm^-1^, that are similar to those reported for canonical Chl *a*-binding PSII under analogous experimental and pre-treatment conditions^**19**^. Figure 3C shows the T-S spectrum recorded upon microwave selection at 998 MHz (│D│+│E│), that displays a main bleaching at 681 nm and an accompanying positive structure between 670 and 677 nm. The presence of both ^3^Chl *d/f* and ^3^Chl *a* in FR-PSII was also confirmed by FDMR (Fluorescence Detected Magnetic Resonance), under conditions of unselective excitation and broadband fluorescence detection at λ>715 nm (Figure S3).

**Figure 3.**
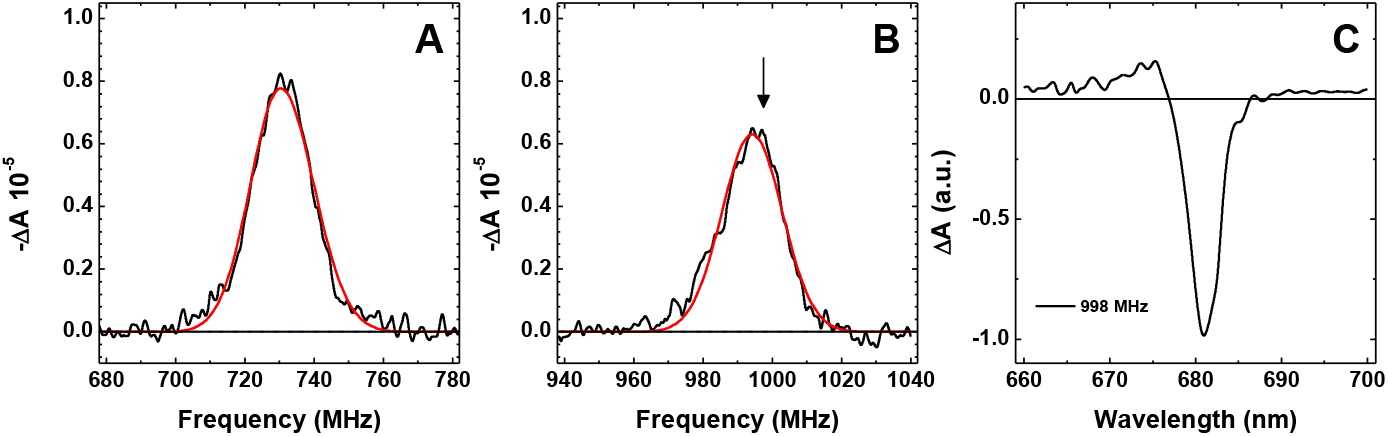
**A** and **B:** ADMR spectra recorded at 682 nm in the ^3^Chl *a* │D│–│E│ and │D│+│E│ transitions respectively (black lines), together with their fit by a single Gaussian band (red line). **C:** T–S spectrum obtained upon microwave selection at 998 MHz (indicated by the arrow in **B**), normalized to its maximal bleaching. Experimental conditions as in Figure 2.

Further insight into the origin of the observed triplet states can be inferred from the population mechanism-dependent esp, that is determinable by TR-EPR. The early time (1.2 – 1.4 μs) TR-EPR spectrum at X-band of FR-PSII under the same reducing conditions employed for the ODMR experiments (Figure 4), confirms the presence of different triplet species. A satisfactorily spectral description is obtained only when considering three triplet components. One is attributed to a carotenoid triplet state that has the broader spectrum and the smallest amplitude. The other two are attributed to ^3^Chl *a* and ^3^Chl *d/f*, based on the respective ZFS parameters (reported together with all simulation parameters and with further detail on the analysis in the SI). Whereas the ^3^Chl *a* is characterized by an *eee/aaa* esp pattern, indicative of its population by intersystem crossing (ISC), the ^3^Chl *d/f* displays an *aee/aae* polarization, characteristic of population by the radical pair recombination mechanism^**14**^. Although the observation of a significantly populated ^3^Chl *a* is both surprising and intriguing, the determined ISC population mechanism excludes its involvement in the photochemical activity of FR-PSII. Its possible origin, perhaps as intermediate of the rChlF catalysis, is discussed in the SI. Further discussion will here focus on the ^3^Chl *d/f*, since its population mechanism links it directly to FR-PSII photochemistry instead.

**Figure 4.**
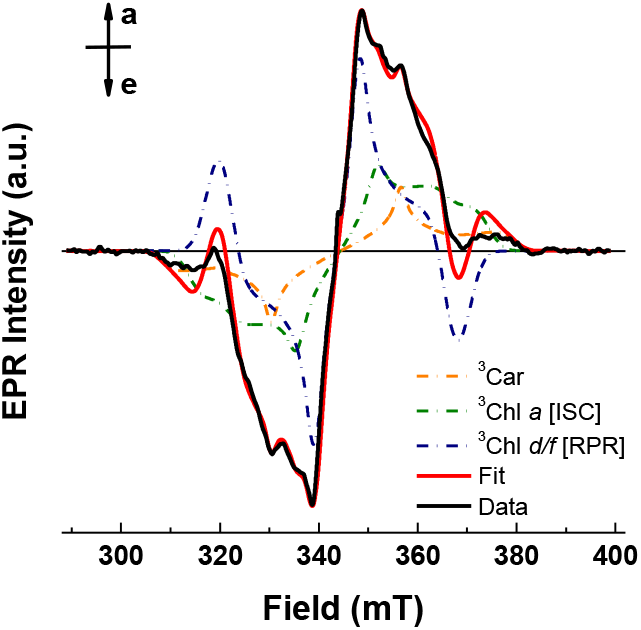
X-band TR-EPR spectrum integrated between 1.2-1.4 μs (black line), together with its simulation (red line) resulting from the contributions of a carotenoid triplet state (orange dash-dotted line), a ^3^Chl *a* with intersystem crossing (ISC) esp (green dash-dotted line) and a ^3^Chl *d/f* with a radical pair recombination (RPR) esp (blue dash-dotted line). Experimental conditions: T = 80 K; υ= 9.65 GHz; mw power: 0.6 mW; excitation wavelength: 532 nm.

Previous studies have identified the Chl_D1_ site as the most probable locus for a red-shifted chlorophyll in FR-adapted PSII RC^**4**,**8-11**^. Consistently, the T–S spectrum in Figure 1B, simulated via diagonalization of an excitonic Hamiltonian with couplings computed under the point-dipole approximation, is supportive of this assignment. Placing the triplet-carrying Chl *d/f* at either the P_D1_ or P_D2_ sites, worsened the spectral simulation (Figure S6). Considering the possible presence of more than one Chl *d/f* molecules in the RC did not lead to any significant improvement of the simulations either (See SI for further detail). These results indicate that Chl_D1_ not only defines the lowest-energy RC state but also represents the energetically preferred site for primary photochemical charge separation and recombination. Low-energy states in the FR-PSII RC are essential to ensure sufficient RC excited-state population under FR illumination, thereby compensating for the simultaneous presence of Chl *d/f* in the CP43 and CP47 antenna complexes. Furthermore, the presence of a *single*, well-defined low-energy state in the RC concentrates the singlet excited state population on the specific site most likely to initiate photochemistry. This represents a different scenario with respect to Chl *a*-only PSII RCs, where the excited state is more evenly delocalized because of the rather contained energy differences of both site energies and excitonic eigenstates^**20**,**21**^. The RC asymmetry brought about by the incorporation of a single Chl *d/f* molecule and the resulting enhanced excited state localization on the putative primary donor, emerges as a key pigment-protein reorganization that preserves both the quantum conversion efficiency of FR-PSII and the water-splitting activity in view of the substantial conservation of the donor-side electron transfer cofactors, P_D1_ and Tyr Z.

## Abbreviations

A/FDMR: Absorption/Fluorescence Detected Magnetic Resonance
Chl: Chlorophyll
(r)ChlF: (recombinant) Chlorophyll *f* synthase
esp: electron spin polarization
FR: Far Red
ISC: Intersystem Crossing
ODMR: Optically Detected Magnetic Resonance
PSII: Photosystem II
RC: Reaction Center
TR-EPR: Time-Resolved Electron Paramagnetic Resonance
ZFS: Zero Field Splitting

## Supplementary information

Additional details concerning biochemical and spectroscopic methodologies, data analysis and spectral simulations. ODMR studies of thylakoid membrane and their analysis. Hypothesis on the origin of the observed Chl *a* triplet.

## Funding Sources

APC, DC and SS acknowledge funding from the EU through MUR, for the project “Extending the red limit of oxygenic photosynthesis: basic principles and implications for future applications”, PRIN20224HJWMH (CUP B53D23015880006/C53D23004620006). AC acknowledges support from the Chemical Complexity C2 project (University of Padova).

## Acknowledgements

We thank Dr. A.W. Rutherford (Imperial College) for the involvement in the preliminary stages of the investigation. The technical assistance of Sabrina Mattiolo (University of Padova) is gratefully acknowledged.

## Notes

### Competing Interest Statement

The authors have declared no competing interest.

