## Supplementary Information for "The Recombination Triplet State in the Far-Red Light Adapted Photosystem II is Located at the Chl_D1_ Site and Resides on the Red-Most Chlorophyll of the Reaction Center"

**Supporting Information: Index.**

1. Supplementary Material and Methods
2. Optical properties of far-red adapted cells, thylakoid membranes and isolated PSII core complexes
3. Extended Scale Triplet minus Singlet (T–S) upon Chl *d/f* microwave selection
4. Fluorescence Detected Magnetic Resonance (FDMR) of Isolated FR-adapted PSII core complex
5. Analysis of the ADMR signals in FR thylakoids and comparison with the FR-PSII core complex
6. Zero field splitting parameters of $^3$Chl *d* and $^3$Chl *f in vitro* and protein-bound
7. Analysis and simulation of the TR-EPR spectrum
8. Simulations of the T-S spectra within the point-dipole approximation.
9. On the origin of the $^3$Chl *a* observed in isolated FR-PSII core complex

### 1. Supplementary Material and Methods

**Biochemical isolation procedures.** Cells of *C. thermalis* PCC7203 were cultivated in BG11 medium in 250 mL cylindrical Drechsel bottles, under continuous humidified-air bubbling (~2 L/min), initially under white light illumination from a LED lamp (3 W, light temperature 2700 K) and then transferred to almost-monochromatic illumination by an array of LED centered at 750 nm (LED750-03AU, Roithner LaserTechnik). After the initial transfer to far-red light, cultures were then maintained and re-inoculated at 750 nm, in order to ensure an as far as possible complete transition to the far-red adapted state. Scale up was performed by increasing the number of cultivation flasks rather than increasing the flask cultivation volume, as this strategy proved more consistent. The cells, upon reaching a scattered-corrected absorption in between 0.8 and 1.2 $cm^{-1}$ at 680 nm (their maximum in the $Q_y$ Chl band), were harvested by centrifugation for 20 min at 5000 RPM and room temperature in a JA-10 Rotor (Beckman). The pellet was then washed two/three times in cold 50 mM HEPES pH 7.5, and further centrifuged at 4 °C, before being frozen in liquid nitrogen until use. Far-red adapted thylakoid membranes (FR-TM) were purified as previously described[1], but an additional thawing/freezing cycle of the cells was performed before breaking and the membranes were further washed in a 50 mM HEPES buffered solution, pH 7.5, to reduce the residual amount of phycobilisomes. FR-TM were frozen in liquid nitrogen and stored at -80 °C. PSII core complexes were isolated essentially as previously described[2] with some modifications. FR-TM were solubilized at a concentration of 400 μg Chl/mL in a 50 mM MES/NaOH (pH 6.5), 10 mM $CaCl_2$, 5 mM $MgCl_2$ buffer with 0.5 % w/v β-dodecylmaltoside for 60 min, under constant mild stirring, on ice. The unsolubilised material was removed by centrifugation (10 min at 16000*x*g, in a refrigerated benchtop centrifuge), and the cleared supernatant loaded on a 0.4-to-1 M sucrose density gradient in 20 mM MES/NaOH (pH 6.5), 10 mM $CaCl_2$, 5 mM $MgCl_2$ buffer with 0.012 % w/v β-dodecylmaltoside, which was run in a SW41Ti rotor (Beckman) for 21 h at 40000 RPM. The higher green-coloured band in the gradient mainly contained the FR-PSII core complexes. Further purification was achieved by ion-exchange chromatography on a Sepharose Q-HP (Amersham) column by a step $MgSO_4$ gradient (from 0 to 100 mM), using the same buffer system but with 0.03 % w/v β-dodecylmaltoside. Each eluted fraction was checked by UV-Vis spectroscopy to ensure sample homogeneity. Only the fractions displaying a consistent absorption spectrum were pooled, further concentrated and desalted by centrifugation-assisted dialysis (10 KDa cut-off). All operations were performed at 4 °C and in darkness or under dim green light. The obtained FR-PSII core complexes at a concentration equivalent to ~ 15 O.D. $cm^{-1}$ at 673 nm were then frozen in liquid nitrogen and stored until use.

**Sample treatment and conditions.** In order to increase the PSII RC triplet yield, the following photoreduction protocol was performed before ODMR and TR-EPR measurements: PSII core complexes were incubated on ice in the dark for 10-15 minutes after the addition of the glucose (10 mM)/ glucose oxidase (~ 1.2 mg/mL)/ catalase (~ 0.24 mg/mL) enzyme mix to remove oxygen[3]. Subsequently, freshly prepared concentrated solutions of sodium dithionite and of PMS (both in a 150 mM HEPES buffer at pH 7.5) were added to the PSII solution. After a dark incubation on ice for 5 minutes, glycerol was added to reach 66 % (*v/v*) concentration. The final dithionite and PMS concentrations (after glycerol addition) were 10 mM and 10 μM, respectively. The resulting PSII solution was then introduced in either the ODMR cell or in the EPR tube. Double-reduction of the quinone acceptors was accomplished by illuminating the sample for 5 minutes at room temperature with a 150 W light projector. Overheating was avoided by ventilating the sample. After illumination, the sample was rapidly transferred in the pre-cooled cryostat at 60 K (for ODMR measurements) or frozen in liquid nitrogen (for EPR measurements).

#### Spectroscopic Methods and Data Analysis

**Steady-state absorption and fluorescence.** Absorption spectra at room temperature were acquired in a Jasco V550 UV-Vis spectrometer, when necessary (cells and thylakoids) adopting scattering-compensation by the opal glasses method. Room temperature fluorescence spectra were recorded in a laboratory-assembled spectrometer, employing a liquid nitrogen cooled CCD camera (LN-CCD, Princeton) coupled to a spectrograph (SpectraPro-300i, Princeton) as previously described[4]. The excitation, which wavelength was set at 435 nm, was provided by 400W Xenon arc lamp filtered by a monochromator (Jasco, J-500A).

**ODMR (FDMR, ADMR and T-S).** ODMR spectra were acquired in a home-built set-up previously described in detail[5], with some modifications. In short, the light from a halogen lamp (250 W, Philips) was focused on the sample cell, which was immersed in an Oxford Spectromag 4 helium bath cryostat (all measurements were carried out at a temperature of 1.8 K), after being filtered through either a 5 cm $CuSO_4$ solution (FDMR spectra) or a 10 cm water filter (ADMR and T-S spectra). In FDMR experiments, the fluorescence was detected through a longpass filter ($\lambda$ >715 nm) using a photodiode, while in absorption-detected experiments, the light transmittance was detected by the same photodiode through a monochromator (Jobin Yvon, mod. HR250). By sweeping the microwave frequency (MW source HP8559b, sweep oscillator equipped with a HP83522a plug-in and amplified by a TWT Sco-Nucletudes mod 10-46-30 amplifier) while detecting the fluorescence/absorption changes at specific wavelengths, the resonance transitions between spin sublevels of the triplet states can be determined. The microwave resonator, where the sample cell is inserted, consisted of a home-built slow pitch (2 mm) copper helix. The microwaves were on/off amplitude modulated for selective amplification and the signal from the detector was demodulated and amplified using a lock-in amplifier (EG&G, mod. 5210). In the updated set-up, the demodulated output from the lock-in amplifier and the photodiode signal was digitized by a NI USB-6003 data acquisition device. The settings of the instrumentation and the signal acquisition parameters were controlled through a laboratory-written code operating in LabView.

**Analysis of FDMR/ADMR spectra.** The spectra were fitted by a laboratory developed software that has been already described in detail[6]. The fit routine employs a linear combination of Gaussian bands as the model function, with the possibility of imposing, when necessary, global constraints to the required fit parameters. The algorithm minimises a $\chi^2$-like maximum likelihood parameter, given by the sum of squared residues weighted by the noise level determined in an off-resonance portion of each spectrum. Minimisation is achieved by a sequential combination of the Simplex algorithm as the initial parameter search and successive refinement by the Levenberg-Marquardt algorithm.

**TR-EPR.** X-band ($\nu$ = 9.65 GHz) TR-EPR spectra were acquired using a Bruker Elexsys E580 spectrophotometer. The sample was introduced in a quartz 3x4 mm (internal and external diameters) tube, subsequently mounted inside a dielectric cavity (Flexline ER 4118X-MD5, Bruker) working in the critically coupled condition. Temperature (set 80 K) was controlled by a liquid nitrogen flow ER 4118 CFO cryostat governed by an iTC Mercury unit (Oxford). Laser excitation was provided by a Quantel Brilliant Nd:YAG pulsed laser equipped with a second harmonic module ($\lambda$ = 532 nm; pulse duration = 10 ns; energy = 2.5 mJ/pulse; repetition frequency = 20 Hz). TR-EPR experiments were performed by measuring the direct transient EPR signal (no lock-in field modulation) in diode detection mode at different magnetic field intensities (110 mT sweep, 0.4 mT step), thus recording a bidimensional data matrix containing the time evolution of the signal at each magnetic field position. Transients were collected, digitized and averaged by an external oscilloscope (LeCroy, mod. 9361) triggered by the laser Q-switch. To extract the reported TR-EPR spectrum, the data matrix was processed by eliminating the intrinsic response of the cavity (by performing a subtraction at an off-resonance magnetic field position far from the sample transitions) and averaging the EPR signal in the 1.2 – 1.4 μs time window after the laser excitation.

**EPR spectra analysis and simulation.** A home-written MATLAB program (version R2024b) based on the EasySpin routine[7,8] library (version 6.0.6) was employed to simulate the powder-like triplet state TR-EPR spectrum. The input parameters, for each triplet species, were the isotropic *g*-factor of each species (2.004), the D and E ZFS dipolar parameters, the triplet sublevel populations and the linewidths. Experimental conditions (such as mw frequency and magnetic field values) were also taken into account. The fit of the experimental spectrum was then obtained by a linear combination of the individual triplet spectra by employing a non-linear least square optimization algorithm managed by the esfit program embedded in EasySpin.

### 2. Optical properties of far-red adapted cells, thylakoid membranes and isolated PSII core complexes

The room temperature absorption spectrum of *C. thermalis* cells adapted to growth on quasi-monochromatic light centered at 750 nm is shown in Figure S1A. In agreement with previous reports (*e.g.* ref. 9) the cells display significant absorption at wavelengths longer than 700 nm and extending beyond 800 nm, consistently with the presence of low-energy absorbing Chl molecules and additional absorption contributions from far-red phycobilisomes (FR-PBS). Nonetheless, the maximal absorption of the cell suspension is found at 678 nm, demonstrating that Chl *a* still represents the main chromophore in the system. Figure S1B shows the room temperature fluorescence emission spectrum of FR-adapted cells, upon excitation at 435 nm under closed PSII centers conditions (induced by the addition of DCMU 20 μM). The maximal emission is found at 748 nm already at room temperature with a clear shoulder at 737 nm. The peak emission is significantly red shifted with respect to organisms in which Chl *a* is the only chlorophyll bound to the photosystem core complexes (*e.g.* ref. 10). Chl *a*-only binding PSII is expected to give rise to an emission band in the 680-685 nm interval. Although some short-wavelength emission is observed in this region, a distinct band could not be clearly identified in the cells, where the integrated fluorescence at $\lambda<700$ nm represents only less than 4% of the integrated emission. This can be taken as an indication that the cells have undergone a transition in which all PSII (and, arguably, PSI) incorporate Chl *d* and especially Chl *f* molecules, acting as the terminal emitters, in the vast majority of the photosystems.

Similar results were obtained by the analysis of the steady-state absorption and fluorescence emission spectra of thylakoid membranes from FR-adapted cells (Figure S1C/D) at RT. The most notable difference in the absorption of FR-adapted cells and FR-TM is, as expected, the marked decrease of the PBS antenna in the ~ 550-650 nm window, as most of the extrinsic light harvesting antenna detaches during TM purification. The emission spectrum of TM shows a maximum at 748 nm, as observed in cells, but the intensity of the 737 nm shoulder is significantly decreased. A weak blue wing emission is also present in TM having a relative maximum at ~683 nm, close to where PSII emission of a standard Chl *a*-binding PSII is expected to be observed. Nonetheless, the integrated emission below 700 nm accounts for only about 3% of the total thylakoid fluorescence.

The steady state absorption and fluorescence emission spectra at RT of the purified FR-PSII core are shown in Figure S1 E/F. The complex displays a maximal absorption at 673 nm. The absorption tail extending in the near-infrared region is clearly visible and, while being less pronounced than in either cells or thylakoids, appears more structured, having a relative maximum at 722 nm and a distinct low-energy shoulder at 741 nm. The narrowing and increased resolution of the long-wavelength tail in the FR-PSII core is due to the disappearance of the overlapped contribution of Chl *d*/*f* absorption forms associated with PSI, in more intact material. The emission of the FR-adapted isolated PSII core shows a maximum at 748 nm and a less pronounced shoulder at 736 nm, at the same wavelengths of cells and TM. Also in the isolated PSII complex, a low intensity short-wavelength emission band is observed, having a broad maximum at 684 nm. The intensity of the fluorescence in the Chl *a* emission window ($\lambda<700$ nm) is however very weak, representing less than 2% of the total integrated emission.

Taken together these observations suggest that, under the growth conditions employed in this investigation, cells have undergone an almost complete transition to FR-adapted photosystems, and that the dominant population of PSII core complexes is represented by those binding low-energy Chl molecules, that give rise to the resolved emission bands centered at 748 nm and 736 nm. The short-wavelength emission ($\lambda<700$ nm) in all samples investigated is always less than 4%, and it is minimal (1.9%) in the FR-PSII core complex. It shall be considered that part of this emission would represent the residual equilibrated fluorescence from Chl *a* in the FR-adapted PSII, since stoichiometrically this remains the most abundant pigment. At the same time, a portion of the short wavelength emission can originate from PSII complexes that have not undergone the FR-acclimation and/or bind primarily Chl *a*. In any case, this would represent a very minor population.

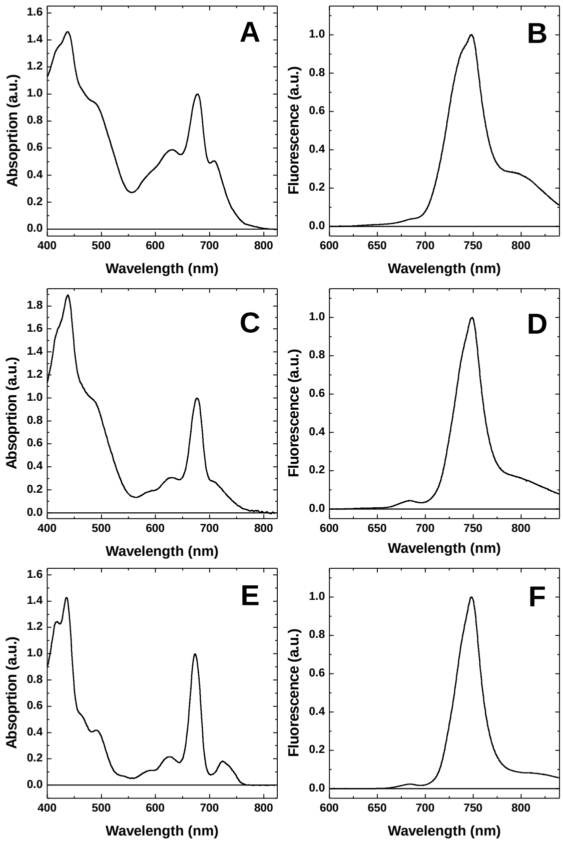


**Figure S1**. The left hand-side panels show the absorption spectra of far-red light (750 nm) adapted cells (**A**), thylakoid membranes (**C**) and PSII core complex (**E**) at room temperature. The absorption spectra are normalized to the maximum in the $Q_y$ Chl *a* transition. The right hand-side panels show the fluorescence emission spectra (excited at 435 nm, FWHM 4 nm) of far-red adapted cells (**B**), thylakoids (**D**) and PSII core complexes (**F**). Fluorescence spectra are normalized to the maximal emission.

#### 3. Extended Scale Triplet minus Singlet (T–S) upon Chl *d/f* microwave selection

Figure S2A shows the T–S spectrum obtained upon 570 MHz microwave selection (|D|–|E| transition) on an extended, non-normalized scale, particularly in the near-infrared spectral region, displaying the triplet-triplet absorption bands. The triplet-triplet absorption extending above 750 nm is relatively featureless, even at the very low temperature (1.8 K) of the ODMR experiments. To the best of our knowledge this spectrum represents the first detection of NIR triplet-triplet absorption in a far-red adapted reaction center, and might therefore provide useful information, particularly for the comparison with differential absorption properties from other species occurring in the same spectral window (e.g. radical pair, chlorophyll cations, etc.)

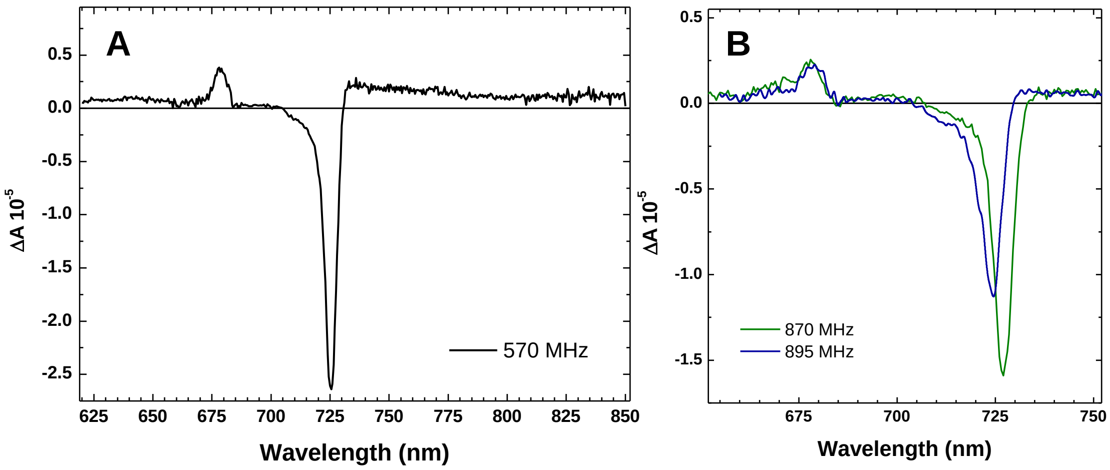


**Figure S2**. **A:** Triplet minus singlet spectrum acquired upon selection at 570 MHz ($^3$Chl *d/f* |D|–|E| transition) showing the extended near-infrared region with respect to the one presented in Figure 1 of the main text. The spectrum also shows the actual ΔA intensity. **B:** Spectra acquired in the |D|+|E| transition region with selection at 870 MHz and 895 MHz showing the actual intensity; the normalized spectra are shown in Figure 2 of the main text. Experimental conditions: T= 1.8 K; Mod. Frequency= 33 Hz; mw power=0.5 W.

In Figure S2B are presented the T–S spectra acquired above selective excitation at 870 MHz and 895 MHz in the |D|+|E| transition, on a non-normalized intensity scale. The absolute intensity of the T–S spectra at the different selection frequencies match closely, as expected, those of the corresponding ADMR spectra (Figure 2).

#### 4. Fluorescence Detected Magnetic Resonance (FDMR) of Isolated FR-adapted PSII core complex

Figure S3 shows the FDMR spectrum acquired upon non-selective broad-band excitation (5 cm $CuSO_4$ 0.6 M + BG12 cut-off filter) and broad-band emission collection (RG715 cut-on filter) that ensure integration over most of the PSII fluorescence spectrum, and hence the recording of all the detectable Chl *a* and *d/f* triplet states. The FDMR displays four clearly distinguished peaks attributable to the |D|–|E| and |D|+|E| transitions of $^3$Chl *a* and $^3$Chl *d/f*. Chl *d/f* triplets fall in the ~500-650 MHz window (|D|–|E|) and the ~800-950 MHz window (D|+|E|), whereas those of Chl *a* in the ~650-800 MHz window (D|–|E|) and the ~950-1050 MHz window (D|+|E|). The position of the FDMR transitions and the overall band-shapes resemble those reported in the main text for the ADMR spectra, associated with $^3$Chl *d/f* upon 725 nm observation (Figure 2) and $^3$Chl *a* upon 682 nm observation (Figure 3). Consistently, the $^3$Chl *d/f* FDMR spectra could be decomposed by a linear combination of two Gaussian sub-bands, having the same center positions (and hence zero-field splitting parameters) as well as the same band-widths used for the ADMR decomposition. Similarly, the $^3$Chl *a* FDMR spectra can be satisfactorily described by a single triplet population. In the case of Chl *a* triplet population, however, it was necessary to slightly shift the band center frequency, by ~ 7 MHz in both transitions, which resulted also in a small variation of ZFS parameters, particularly the value of |D| varied from 0.0288 $cm^{-1}$ (ADMR) to 0.0290 $cm^{-1}$ (FDMR), as reported in **Table S1**. These differences may be due to the higher selectivity of the ADMR spectra, detected at specific wavelengths, compared to the FDMR spectra which result from the integrated fluorescence spectrum.

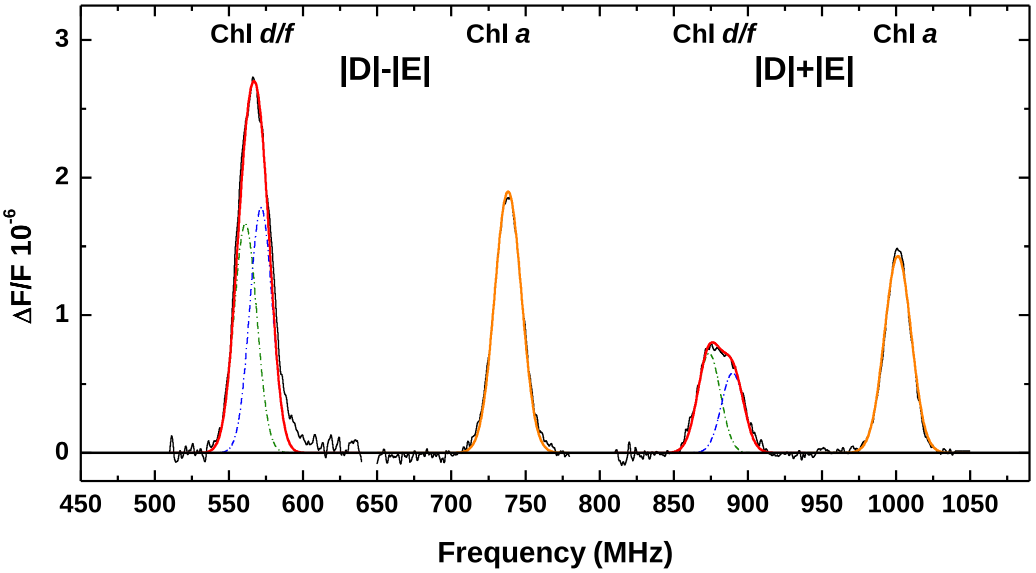


**Figure S3**. FDMR spectrum of isolated far-red adapted PSII core complex, obtained from broad-band excitation (λ< 500 nm) and unselective detection (λ> 715 nm), integrating most of the photosystem emission. The experimental traces are shown as black lines, and the corresponding transitions of specific Chl molecules are indicated in the figure. Also shown is the decomposition using a linear combination of Gaussian bands of the $^3$Chl *d* |D|–|E| and |D|+|E| (dash-dotted blue and green lines: sub-bands; red line: fit), and the description by a single Gaussian band (orange lines) of the $^3$Chl *a* |D|–|E| and |D|+|E| transitions. Experimental conditions: T= 1.8 K; Mod. Frequency= 33 Hz; mw power=0.5 W.

**Table S1. Fit Parameters describing the FDMR and ADMR spectra in the far-red adapted PSII core complex**

| \|D\|–\|E\| (MHz) | \|D\|+\|E\| (MHz) | FWHM (MHz) | \|D\| ($cm^{-1}$) | \|E\| ($cm^{-1}$) | Assignment |
|---|---|---|---|---|---|
| $^3$Chl *d/f* | | | | | |
| 561.1 ±0.5 | 873.8 ±0.5 | 18.0 ±0.5 | 0.0239 | 0.0052 | $^3$Chl *d/f* FR-PSII RC (ADMR/FDMR) |
| 571.6 ±0.5 | 889.7 ±0.5 | 17.7 ±0.5 | 0.0244 | 0.0053 | $^3$Chl *d/f* FR-PSII RC (ADMR/FDMR) |
| $^3$Chl *a* | | | | | |
| 730.4 ±0.5 | 994.2 ±0.5 | 21.4 ±0.5 | 0.0288 | 0.0044 | $^3$Chl *a* FR-PSII Core/ChlF ADMR |
| 738.3 ±0.5 | 1001.2 ±0.5 | 21.3 ±0.5 | 0.0290 | 0.0044 | $^3$Chl *a* FR-PSII Core/ChlF FDMR |

The table reports the parameters retrieved from the constrained fit of the FDMR (Figure S3) and ADMR (Figures 2 and 3) spectra, and the resulting zero-field splitting parameters |D| and |E| for the different $^3$Chl *d*/*f* and $^3$Chl *a* populations.

### 5. Analysis of the ADMR signals in FR thylakoids and comparison with the FR-PSII core complex

Panels A and B of Figure S4 show a direct comparison of the ADMR spectra recorded at an observation wavelength of 725 nm in the thylakoid membranes and the PSII complex isolated from FR-adapted *C. thermalis* cells, and subjected to the same pre-illumination under reducing conditions before the measurement. The spectra have been each normalized in the |D|–|E| transition and the same scaling factor is applied to the |D|+|E| to conserve the respective intensity ratios. The comparison demonstrates that the signals acquired in the isolated PSII complexes are very similar to those observable in the TM, including the occurrence of two readily discernible $^{3}$Chl *d/f* sub-populations especially in the |D|+|E| transition. The only notable difference between the FR-TM and FR-PSII ADMR spectra recorded at 725 nm is a variation in the overall band-shape that can be however explained, as shown in the Gaussian sub-band decomposition reported in Figure S4C/D, by changes in the relative intensities of the two $^{3}$Chl *d/f* populations. The population having ZFS |D|=0.0244 cm$^{-1}$, |E|=0.0053 cm$^{-1}$ is slightly more represented than the one characterised by |D|=0.0239 cm$^{-1}$, |E|=0.0052 cm$^{-1}$ in TM, whereas the opposite partition is determined in the isolated PSII core complex. It is worth stressing, that the fits of the ADMR spectra (725 nm) in FR-TM were obtained with the exact same center frequencies and bandwidths employed for the isolated FR-PSII to describe the ADMR spectra (Table S1). Since TM can be considered a relatively intact material that is, moreover, purified without the aid of detergents, it can be safely excluded that the occurrence of two $^{3}$Chl *d/f* populations associated to the RC of PSII complex represent an artefact, rather these reflect a characteristic feature of the FR-adapted photosystem.

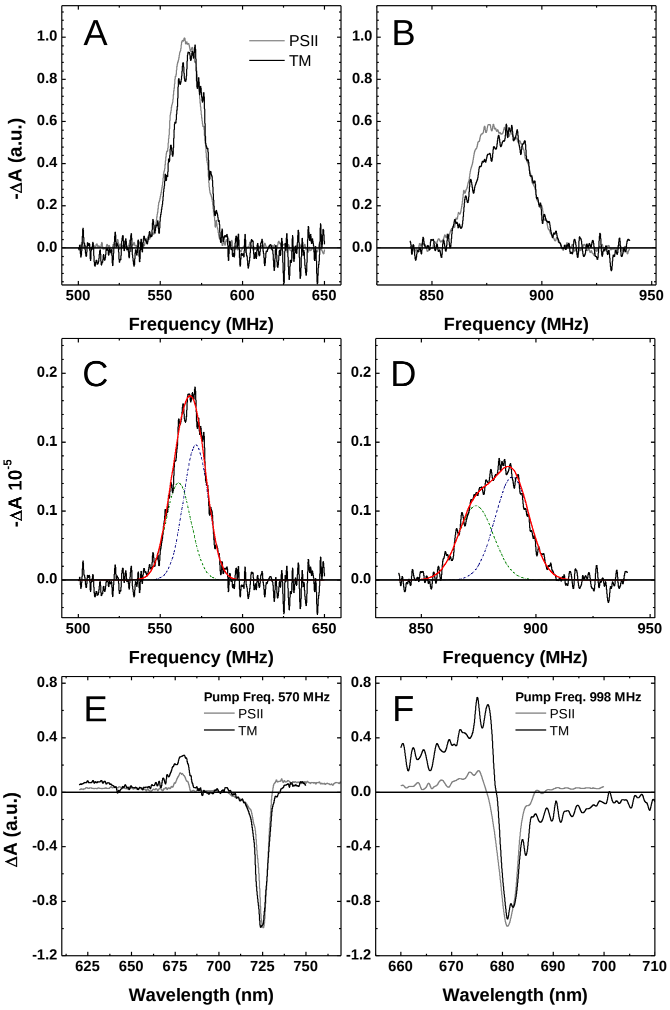


**Figure S4**. ADMR spectra recorded at 725 nm in far-red adapted thylakoid membranes (black lines) and PSII core complexes (gray lines) in the $^{3}$Chl *d/f* |D|–|E| (**A**) and |D|+|E| (**B**) transitions. Note that the spectra were normalised in the |D|–|E| (and equally scaled in the |D|+|E|) to allow for direct comparison. **Panels C/D** show the decomposition of the TM ADMR spectra using a linear combination of Gaussian bands (dash-dotted blue and green lines: sub-bands; red line: fit) in the |D|–|E| (**C**) and |D|+|E| (**D**) transitions of $^{3}$Chl *d/f*. The fit parameters are, except for the amplitudes, those reported in Table S1. Panel **E**: mw-induced T–S spectra (570 MHz, $^{3}$Chl *d/f*

|D|–|E|) recorded in FR-TM (black line) and FR-PSII complex (grey line). Panel **F**: mw-induced T–S spectra (998 MHz, $^3$Chl *a* |D|+|E|) in FR-TM (black line) and FR-PSII complex (grey line). Each spectrum is normalised to its ground state bleaching maximum. Experimental conditions as in the legend of Figure 2 and S3.

The robustness of the data obtained in FR-PSII is further confirmed by the comparison of the T–S spectra recorded upon mw selection at 570 MHz (|D|–|E|) in the isolated core complex and TM, shown in Figure S4E. The lower signal-to-noise ratio in TM is to be expected because of both the intrinsic scattering nature of the membranes and the contribution of both PSI and PSII absorption to the overall TM spectrum, so that the relatively small absorption changes induced by the mw modulation need to be recorded over a larger background. The latter can also explain the relatively contained difference in the intensity of the main spectral features, that, nonetheless, are detected at the same wavelengths. The main ground state bleaching falls at 724.5 nm and the coupled side excitonic bands between ~660 and 690 nm.

ADMR spectra at 680 nm, which showed a clear contribution from a $^3$Chl *a* in FR-PSII (Figure 3), fell below the noise level in thylakoids. It was nonetheless possible to record a T– S spectrum upon mw selection at 998 MHz (in the $^3$Chl *a* |D|+|E| transition). This spectrum is also compared to the one obtained in purified FR-PSII upon the same frequency selection (Figure S4F). The 998 MHz-induced T– S spectra of FR-TM and FR-PSII show main singlet bleaching at around 680 nm and positive, less structured, counter-band centered at ~675 nm, however the intensity of the two main spectral features is different in the two samples. As already mentioned for the 570 MHz-selected T– S spectrum, the most likely reason is the occurrence of some spectral distortion (principally due to so-called "flattening") in TM, which would be even more severe for the 998 MHz-induced ($^3$Chl *a*) since most of the spectral differences are overlapped with the maximal absorption of the sample, whereas at least the main bleaching occurs outside the maximal absorption for the 570 MHz ($^3$Chl *d/f*) selection.

It is also interesting to notice that the identified resonance frequencies of Chl *d/f* represent specific FR-PSII spectroscopic markers, since (FR)-PSI which is also present in the thylakoid membrane, does not contribute to any significant extent to either the ADMR (S4A-D) or the T–S spectrum (S4E).

### 6. Zero field splitting parameters of $^3$Chl *d* and $^3$Chl *f in vitro* and protein-bound

A compilation of the ZFS parameters, $\lvert D\rvert$ and $\lvert E\rvert$, of $^3$Chl *d* and $^3$Chl *f* resulting from ODMR and TR-EPR investigations at cryogenic temperatures is presented in Table S2. When possible, a comparison of values retrieved from the two methods in the same environment (either organic solvents or protein-bound) is shown. Table S2 also show a selection of $^3$Chl *a* ZFS parameters retrieved from ODMR studies of oxygenic RC. The table reports the $\lvert D\rvert-\lvert E\rvert$ and $\lvert D\rvert+\lvert E\rvert$ frequency regions (in MHz) to facilitate the comparison with the experimental ADMR and FDMR spectra reported in this study.

**Table S2. Comparison of the zero field splitting parameters of $^3$Chl *d*, $^3$Chl *f* and $^3$Chl *a***

| $\lvert D\rvert-\lvert E\rvert$ (MHz) | $\lvert D\rvert+\lvert E\rvert$ (MHz) | $\lvert D\rvert$ ($cm^{-1}$) | $\lvert E\rvert$ ($cm^{-1}$) | Detection | Environment |
|---|---|---|---|---|---|
| $^3$Chl *d* | | | | | |
| Organic solvents | | | | | |
| 599 | 887 | 0.0247 | 0.0048 | ODMR | MeTHF[11] |
| 627 | 926 | 0.0259 | 0.0050 | TR-EPR | MeTHF[11] |
| Protein-bound (RC) | | | | | |
| 612 | 861 | 0.0246 | 0.0042 | ODMR | $^3P_{740}$ *A. marina* PSI[12] |
| 629 | 861 | 0.0248 | 0.0039 | ODMR | $^3P_{740}$ *A. marina* PSI[12] |
| 608 | 858 | 0.0245 | 0.0042 | ODMR | $^3P_{740}$ *A. marina* TM[13] |
| 623 | 864 | 0.0248 | 0.0040 | ODMR | $^3P_{740}$ *A. marina* TM[13] |
| 597 | 860 | 0.0243 | 0.0044 | TR-EPR | $^3P_{740}$ *A.* marina PSI[13] |
| $^3$Chl *f* | | | | | |
| Protein-bound | | | | | |
| 600 | 905 | 0.0251 | 0.0051 | ODMR | rChlF synthase[14] |
| $^3$Chl *a* | | | | | |
| Protein-bound (RC) | | | | | |
| 721 | 991 | 0.02855 | 0.0045 | ODMR | $^3P_{680}$ BBY[15] |
| 730-740 | 970-992 | 0.02855, 0.02888 | 0.00388, 0.00422 | ODMR | $^3P_{680}$ D1/D2/$cyt_{559}$[16,17] |
| 720 | 991 | 0.0285 | 0.0045 | ODMR | $^3P_{680}$ TM[18] |
| 715 | 942 | 0.0276 | 0.00379 | ODMR | $^3P_{700}$[19,20] |
| 732 | 952 | 0.0281 | 0.00367 | ODMR | $^3P_{700}$[19,20] |
| 720 | 942 | 0.0277 | 0.0037 | ODMR | $^3P_{700}$ TM[18] |
| 733 | 951 | 0.0281 | 0.0036 | ODMR | $^3P_{700}$ TM[18] |

### 7. Analysis and simulation of the TR-EPR spectrum

The experimentally recorded TR-EPR spectrum presented in Figure 4 of the main text shows a complex pattern that cannot be satisfactorily simulated when considering the contribution from just a single triplet species. This observation is in agreement with the ADMR/FDMR results, which demonstrate the presence of both Chl *d/f* and of Chl *a* triplet states. Whereas in zero-field ODMR the transitions of these two species are well separated (on the mw frequency axis), they are still distinguishable but overlapped in the presence of an external magnetic field. On top of the triplet originating from the two types of Chls, including the contribution of a broader component which, based in its characteristics[14,21], can be assigned to a carotenoid triplet ($^3$Car), was also necessary to obtain a satisfactory agreement between the experimental and the simulated TR-EPR spectrum (Figure 4). The main parameters obtained from the analysis and simulation of the TR-EPR spectrum are reported in Table S3 below.

**Table S3. Parameters employed in the fit of experimental and simulations of TR-EPR spectra**

| | Zero Field Splitting | | | | Zero field populations (High field populations) | | | Isotropic Linewidth | H Strain | | |
|---|---|---|---|---|---|---|---|---|---|---|---|
| | \|D\| (MHz) | \|E\| (MHz) | D ($cm^{-1}$) | E ($cm^{-1}$) | $p_X/(p_{-1})$ | $p_Y/(p_0)$ | $p_z/(p_{+1})$ | (mT) | $H_X$ (mT) | $H_Y$ (mT) | Hz (mT) |
| $^3$Chl *a* | 848 | 131 | 0.0283 | -0.0044 | 0.39 | 0.43 | 0.18 | 2.5 | 4.2 | 0.3 | 4.6 |
| $^3$Chl *d/f* | 738 | 161 | 0.0246 | -0.0054 | (0) | (1) | (0) | 2.5 | 4.2 | 0.3 | 4.6 |
| $^3$Car | 1000 | 89 | -0.0333 | -0.0030 | 0.30 | 0.33 | 0.37 | 1.5 | 0.2 | 0.4 | 4.1 |

The table lists the parameters extracted from the best fit of the experimental TR-EPR spectrum, considering triplets from three chromophores: a $^3$Chl *a* populated by intersystem crossing (ISC), a $^3$Chl *d/f* populated by the radical pair recombination mechanism (RPR) and a $^3$Car. The results of the fitting in terms of ZFS values (D and E), triplet sublevel populations and line-widths for each of the species considered are all summarized. The microwave frequency in the simulations was 9.65 GHz and the isotropic *g*-factor was 2.004 for all species (dipolar interactions give a substantial broadening of the EPR spectrum thus making *g*-factor anisotropies negligible).

The key difference evidenced from the analysis is that, whereas the $^3$Chl *a* is characterized by a polarization pattern stemming from intersystem crossing (ISC) population mechanism, the $^3$Chl *d/f* shows a polarization pattern that is unambiguously associated to the radical pair recombination (RPR) mechanism, and has therefore to be originated in the FR-PSII reaction center. Figure S5 shows the spectra of the simulated species, each normalized to the same intensity to better highlight their spectral differences. Moreover, for the $^3$Chl *d/f* species, on top of the retrieved spectrum showing the *aee/aae* pattern resulting from the RPR mechanism, the spectrum which would be observed for an ISC population mechanism, is also shown. Similarly, the comparison between the retrieved ISC-populated $^3$Chl *a* and the one which will be expected from the RPR mechanism are directly compared.

It should be appreciated that not only the spectra of the ISC- and RPR-populated triplets are clearly distinguished, but also it would not be possible to reproduce the experimental spectrum by assuming an opposite situation that comprises an ISC-populated $^3$Chl *d/f* and an RPR-populated $^3$Chl *a*, supporting the suggestion that they originate from Chl-binding complexes having different composition and photochemical activity. It shall also be noted that this finding seems to exclude the possibility that any sizable population of non FR-adapted PSII centers, binding only Chl *a* in both the RC and the core complexes, is present in the sample. This population would give rise to $^3$Chl *a* displaying the *aee/aae* spin polarization[22,23], which is not observed instead. Concerning the $^3$Car detected, the simulation is fully compatible (in dipolar D and E parameters and triplet sublevel populations) with the one obtained from TR-EPR measurements on the recombinant rChlF enzyme[14].

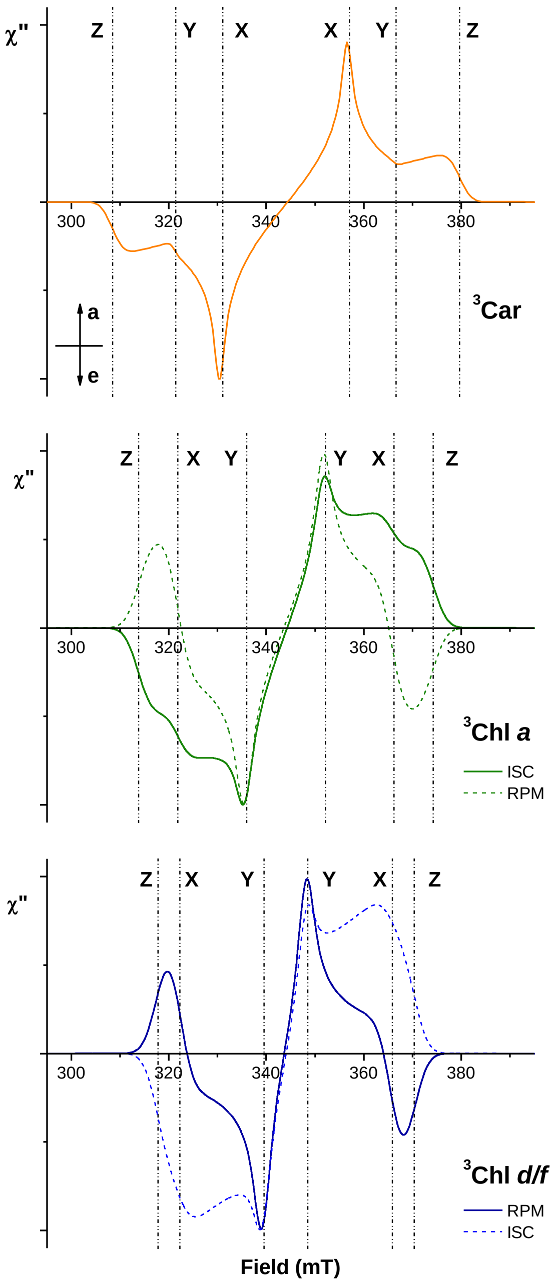


**Figure S5**. Simulated TR-EPR spectra of the isolated triplet components obtained from the decomposition of the experimental spectrum shown in Figure 4 of the main text. The simulated $^3$Car (orange lines), $^3$Chl *a* (green lines) and $^3$Chl *d/f* (blue lines), scaled to the same amplitude and showing the canonical axis transitions, are reported independently. For the Chl triplets are also shown the spectra expected for the complementary population mechanism. Hence, for $^3$Chl *a* the experimentally retrieved ISC-populated spectrum (solid line) is compared to the RPR-populated simulation (dashed line). For $^3$Chl *d/f*, the experimentally retrieved RPR-mechanism spectrum (solid line) is compared with the potential ISC-populated triplet simulation (dashes line).

### 8. Simulations of the T–S spectra within the point-dipole approximation

**Theory and spectra simulations**. To simulate the T–S spectrum of the RC recombination triplet state, an excitonic Hamiltonian matrix ($\widehat{H}_{ex}$), describing the system's energetics, was established:

$$\widehat{H}_{ex} = \sum\nolimits_{a} E_a |a\rangle\langle a| + \sum\nolimits_{a\neq b} V_{ab}|a\rangle\langle b|$$

where $E_a$ is the site energy of pigment $a$ (defined as the vertical transition energy of the pigment, keeping the equilibrium position of the nuclei of the electronic ground state) and $V_{ab}$ is the coupling term between each pair of pigments $a$ and $b$[24].

The eigenvalue problem to be solved (given by the diagonalization of the exitonic matrix) is then $\widehat{H}_{ex}\boldsymbol{\Psi}_m = E_m\boldsymbol{\Psi}_m$, where $E_m$ is the energy of each $\boldsymbol{\Psi}_m$ exitonic state. In the pigment representation, every $\boldsymbol{\Psi}_m$ eigenvector can be expressed as the linear combination $\boldsymbol{\Psi}_m = (c_1^m, c_2^m, \ldots, c_n^m)$ where $c_n^m$ is the participation of the $n$-th pigment in the $m$-th exitonic state. The absorption spectrum of the collection of pigments is estimated as $Abs \propto \sum_m |\boldsymbol{\mu}_m|^2$, where $\boldsymbol{\mu}_m$ is the transition dipole moment (TDM) of the $m$-th exciton, calculated as $\boldsymbol{\mu}_m = \sum_n c_n^m \boldsymbol{\mu}_n$, with $\boldsymbol{\mu}_n$ being the TDM of the $n$-th pigment.

The interaction term $V_{ab}$ was estimated by adopting the simple point-dipole approximation, which describes the electrostatic interaction between each couple of pigments as a point dipole - point dipole interaction of the TDMs associated to the single molecules composing the collection of pigments. The off-diagonal elements of the exitonic Hamiltonian matrix are then $V_{ab} = \frac{1}{4\pi\varepsilon_0}\kappa_{ab}\frac{\mu_a\mu_b}{R_{ab}^3}$ where $\mu_a$ and $\mu_b$ are the TDM intensities of the interacting couple of pigments $a$ and $b$, and $R_{ab}$ their distance. The relative orientation between TDMs is taken into account by the orientation factor defined as: $\kappa_{ab} = \widehat{\boldsymbol{\mu}}_a \cdot \widehat{\boldsymbol{\mu}}_b - 3(\widehat{\boldsymbol{\mu}}_a \cdot \widehat{\boldsymbol{R}}_{ab})(\widehat{\boldsymbol{\mu}}_b \cdot \widehat{\boldsymbol{R}}_{ab})$, where $\widehat{\boldsymbol{\mu}}_a$ and $\widehat{\boldsymbol{\mu}}_b$ are the TDM versors and $\widehat{\boldsymbol{R}}_{ab}$ the versor of $R_{ab}$.

The position and mutual orientation of the TDM of each pigment considered in the exciton model, obtained from atomic coordinates, were initially retrieved from the high-resolution crystal-structure of PSII (PDB: 3WU2[25]). The coordinates from a FR-adapted structural model (PDB 8EQM[26]) were also considered for comparison. TDMs were considered to be oriented along the segment connecting the $N_B$-$N_D$ nitrogen atoms of the chlorin ring. Distances between pigments were determined as the distance between the centers of mass of the $N_B$ and $N_D$ atoms.

Transition dipole moments for the different pigments were taken as 4.4 D[27], 3.5 D[27] and 5.0 D[28] for respectively Chl *a*, Pheo *a* and Chl *d/f.* The difference in the estimated extinction coefficient of the intrinsic long-energy Chl *d* and Chl *f* molecules were estimated to be relatively similar[29]. Hence, the effective transition dipole of 5.0 D shall be representative of both chromophores. These TDMs have to be considered as *effective,* thus already taking into account the dielectric effects of the protein medium[30,31].

Since the triplet is a neutral species, the T–S spectra are dominated by exciton redistribution. The effect of local electric charges on the site energies is not negligible in the description of radical- and radical pair-induced absorption differences instead. This in turn significantly simplifies the T–S spectral simulation which can be reasonably reproduced considering a minimal set of variables, being substantially only the unperturbed site energies of the chromophores (the diagonal elements of $\widehat{H}_{ex}$) and the chromophore on which the triplet state is localized. The T–S spectrum is then simulated, by initially solving the excitonic Hamiltonian, that comprised the 8 Chl pigments coordinated by the D1/D2 heterodimer. This provides the description of the singlet excited states of the system. Subsequently, the pigment where the triplet state is supposed to be located (for example, at the $Chl_{D1}$ position) is removed from the exciton system and the calculation is repeated. Hence, the description of the system in its triplet state is provided by a solution of the Hamiltonian with a $n$-1 dimension, and all other parameters are the same as of the singlet representation. An example of the exciton matrix to be diagonalized, for the system fully in its singlet state , is reported in Table S4.

| | $P_{D1}$ | $Chl_{D1}$ | $Pheo_{D1}$ | $P_{D2}$ | $Chl_{D2}$ | $Pheo_{D2}$ | $ChlZ_{D1}$ | $ChlZ_{D2}$ |
|---|---|---|---|---|---|---|---|---|
| $P_{D1}$ | 15020 | -20.864 | -3.69 | 243.195 | -87.799 | 17.146 | 0.551 | 1.018 |
| $Chl_{D1}$ | -20.864 | 13800 | 89.964 | -97.475 | 15.897 | -6.017 | 3.19 | -0.077 |
| $Pheo_{D1}$ | -3.69 | 89.964 | 14840 | 19.754 | -5.822 | 3.191 | -3.644 | -0.289 |
| $P_{D2}$ | 243.195 | -97.475 | 19.754 | 15020 | -12.067 | -4.3 | 1.155 | 0.928 |
| $Chl_{D2}$ | -87.799 | 15.897 | -5.822 | -12.067 | 14990 | 72.275 | -0.137 | 2.447 |
| $Pheo_{D2}$ | 17.146 | -6.017 | 3.191 | -4.3 | 72.275 | 14790 | -0.288 | -3.83 |
| $ChlZ_{D1}$ | 0.551 | 3.19 | -3.644 | 1.155 | -0.137 | -0.288 | 14990 | 0.221 |
| $ChlZ_{D2}$ | 1.018 | -0.077 | -0.289 | 0.928 | 2.447 | -3.83 | 0.221 | 14970 |

**Table S4.** Hamiltonian matrix (in $cm^{-1}$) describing the excitonic system of the far-red PSII. The considered pigments were 3 Chls *a* (RC), 1 Chl *d/f* (RC), 2 Pheo *a* (RC) and the two so-called $Chl_Z$ molecules, at the periphery of the complex. Diagonal elements represent the site energies, the off-diagonal elements represent the pigment-pigment interactions, estimated within the point-dipole approximation.

In order to obtain a simulated spectrum whose bandshape resembles the experimental one and hence providing a meaningful comparison, each excitonic state was dressed using a Gaussian band centered at the exciton transition energy ($E_m$) and having an area equivalent to the exciton transition intensity ($|\boldsymbol{\mu}_m|^2$). The widths of all Gaussian bands were set to a FWHM of 113 $cm^{-1}$. The excitonic transitions in the singlet and triplet state of the system were dressed independently and then subtracted to yield the simulated T–S spectrum. No attempt was made to describe Triplet-Triplet absorption features that, in the investigated spectral window, occur at wavelength longer that ~ 740 nm. All calculations were performed using a laboratory-written MATLAB (R2024b version) script.

**Discussion of the impact on the simulation of the pigment site energies and triplet-carrying sites.** The most satisfactorily description of the experimental T–S spectrum was obtained when considering for the system in its singlet state, site energies almost identical to those already suggested by Renger and coworkers[30] (reported in the structure of Figure 1A), and the triplet state sitting in the $Chl_{D1}$ position, representing the red-most transition in the system. The only minor modifications to unperturbed site energies were a red shift of 2 nm for $Pheo_{D1}$ and of 1 nm for $Pheo_{D2}$. The site energy of low-energy Chl *d/f* was set at 724.5 nm, which basically coincides with the experimental main ground state bleaching (parameters are reported in Table S4).

It is worth stressing that the main scope of these simulations was to establish the site in which the triplet state resides rather than to obtain a detailed description of the system energetics, which will require more in-depth investigations and more sophisticate approaches for the estimation of both excitonic couplings and unperturbed site energies. Nevertheless, the point-dipole approximation provides a sensible description and allows for a simple comparison of simulations in which the triplet state sits on different sites of the system.

Figure S6 shows the comparison of simulated T–S spectra setting the lower energy state, coinciding with the triplet-carrying state, at the different pigments within the FR-PSII reaction center, *i.e.* $P_{D1}$, $P_{D2}$ and $Chl_{D2}$. The simulations indicated that positioning the low energy state, and the triplet, at $P_{D1}$ or $P_{D2}$ (site energy 721 nm, in both cases) positions, worsen the quality of the simulations, as an additional structure in the high energy exciton band around 655-675 nm is predicted by the simulations, but is not observed experimentally.

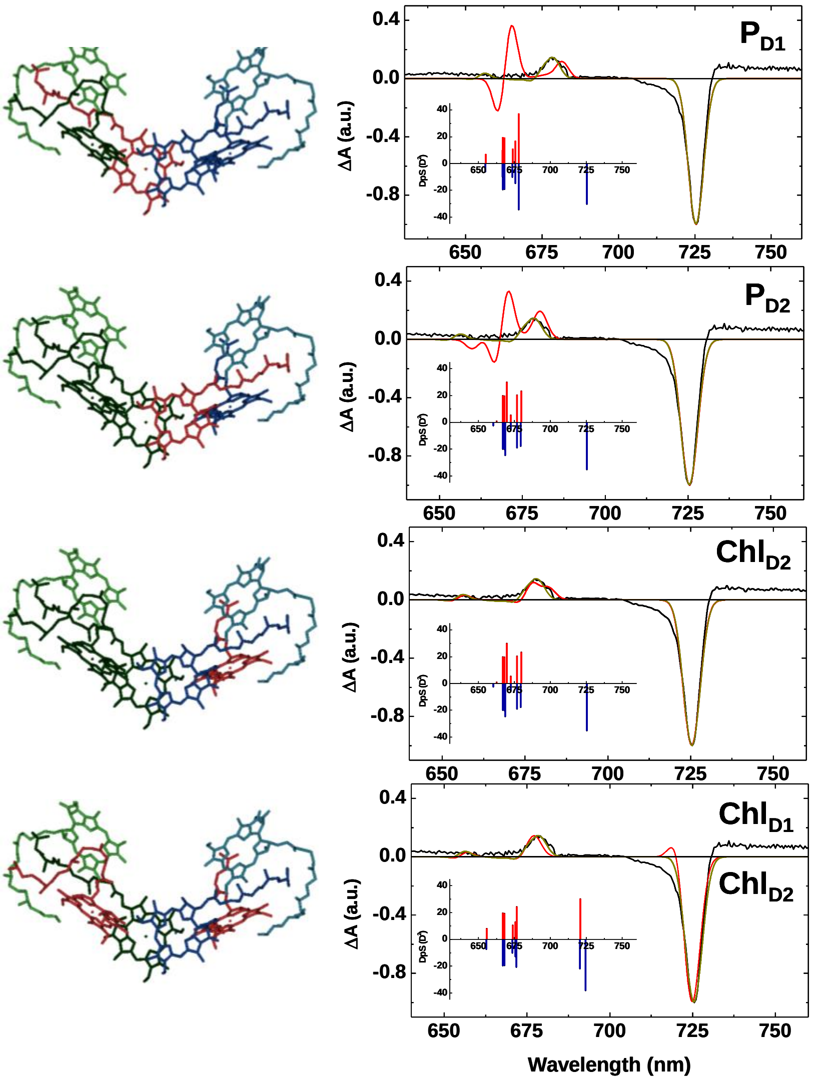


**Figure S6.** Comparison of the simulated (red lines) and experimental (black lines) T–S spectra (570 MHz selection) when considering the low-energy and triplet carrying state sitting on $P_{D1}$, $P_{D2}$, Chl$_{D2}$ and Chl$_{D1}$-and-$_{D2}$, as indicated by the red-colored chromophores in the left-hand side cartoons. The T–S spectrum obtained for the triplet sitting on Chl$_{D1}$ is shown in each panel (gold lines), for ease of comparison. The insets in the right-hand panels show the stick-spectra representing the eigenstates and corresponding population (in $D^2$ units) in the singlet (blue bars, negative sign set arbitrarily) and triplet (red bars, positive sign) for each simulation. All spectra are normalized to their maximal ground state bleaching.

In the case of the low-energy state at the Chl$_{D2}$ (site energy 724.5 nm as for Chl$_{D1}$) position, the simulation is overall more comparable to the one obtained for the Chl$_{D1}$ position (see also the main text), but for slight broadening of the (positive) excitonic feature, which might not be that significant within the adopted approximation. Nonetheless, the presence of a low energy state on a site within the inactive electron transfer chain appears inconsistent with the functionality of the system and with the triplet population by the radical pair recombination mechanism.

Further, Figure S6 shows the simulations in which two low energy states are included in the system, at both Chl$_{D1}$ and Chl$_{D2}$ positions (site energies 723.5 nm and 720 nm, respectively), with the triplet sitting on the former. In this case quality of the simulation worsens slightly (appearing of a positive side band feature next to the main bleaching, and narrowing of the high energy excitonic band) rather than improving. In particular, considering a second low energy state does not compensate for the discrepancy between measured and simulated spectra around the main ground state bleaching (the broadening centered at ~713 nm), supporting the notion that a single low energy state is sufficient to account for the experimental observations.

To confirm the robustness of the assignment to the chromophore in the Chl$_{D1}$ as the low-energy (Chl *d*/*f*) and triplet–carrying state, the excitonic calculations were performed also by considering an isoenergetic system, where all Chl *a* and Pheo *a* site energies were set at 666 nm, as well as by performing the calculation considering the coordinates from the structural model of a far-red adapted PSII isolated from *Synechococcus sp.* PCC7335[26] (PDB 8EQM) that also suggested the presence of a low energy element, assigned to Chl *d*, in the $Chl_{D1}$ site.

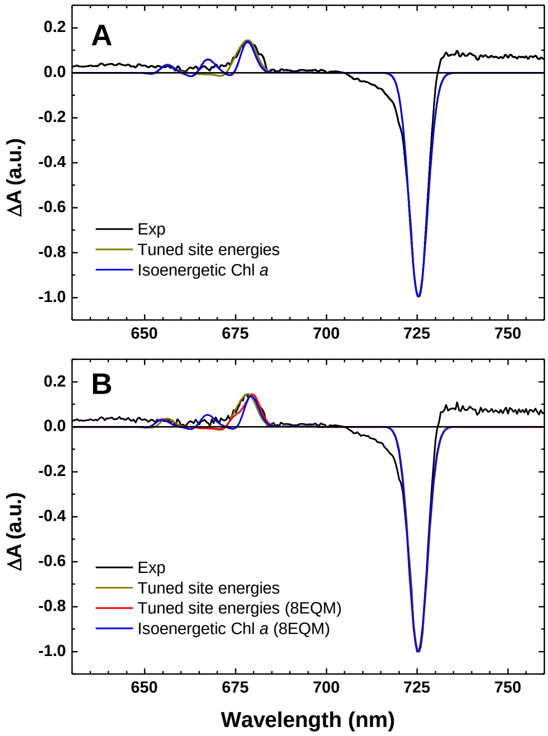


**Figure S7. A:** Comparison of the experimental (black lines) T–S spectra (570 MHz selection) and those simulated considering tuned site energetics (green lines) and isoenergetic site energies (666 nm, blue lines) for all pigments other than $Chl_{D1}$ (724.5 nm) where the triplet, when populated resides, employing the coordinates from the 3WU2 PDB. **B**: Comparison of the experimental (black lines), reference simulation with tuned energies (3WU2 PDB coordinates) and same tuned (red line) and isoenergetic (666 nm, blue lines) site energies but employing the coordinates from the FR-PSII (PDB 8EQM) structural model.

Figure S7 shows that in both cases, the main features of the experimental spectrum are substantially well reproduced. As expected, for the case of an isoenergetic system (excluding the $Chl_{D1}$ site), there are some, but not large, discrepancies in the high energy exciton band. These are in any case less pronounced that what simulated for a low energy state sitting on either $P_{D1}$ or $P_{D2}$. For the simulations resulting from the FR-PSII coordinates, even by using the same site energies with respect to those of *T. vulcanus* (non-far-red adapted PSII; PDB 3WU2[25]) a satisfactorily description was readily obtained. Also in this case, when considering isoenergetic sites for the non-low energy pigments, the quality of the simulation worsened but not in a dramatic fashion

Taken together, these observations indicated that, although more detailed calculations will almost certainly be necessary to provide a refined description of the site energies in the reaction center of the FR-adapted PSII, the identification of low-energy state, on which the recombination-populated triplet state resides, can be considered as reliable.

### 9. On the origin of the $^3$Chl *a* observed in isolated FR-PSII core complex

The analysis of both ODMR (Figures 3, S3 and S4) and TR-EPR (Figure 4) demonstrates the presence of a photoinduced $^3$Chl *a* triplet in the FR-PSII core complex preparation. Similar signals could also be recorded by ADMR in FR-TM that shows a T–S spectrum, upon the same microwave selection employed for purified PSII, having an overall similar bandshape, considering the potential spectral distortion in a scattering material. Hence, the observed $^3$Chl *a* does not appear to be due to possible purification artefact, since TM are a rather intact environment. Although exact quantification of the ODMR signal in terms of triplet yield is not possible, because of differences in polarization and differences in response of the resonator in non-saturating conditions, still, the $^3$Chl *a* population is clearly detected and deserves further discussion.

It is worth noting that the T–S spectrum associated with the $^3$Chl *a* triplet in FR-PSII closely resembles that of $P_{680}$ in the canonical PSII reaction center, whereas a similar spectral signatures are not observed for triplet states associated with light-harvesting complexes.

The simplest explanation would be that the observed $^3$Chl *a* resides in a population of PSII complexes that has not undergone the FR acclimation, and is therefore, substantially, a canonical Chl *a*-binding PSII. Several observations argue against this interpretation. In the first place, the intensity of the detected ODMR is far larger than what would be expected on the basis of the short-wavelength fluorescence emission which could be attributed to Chl *a*, and was estimated to be in between 2-5% both in the FR-PSII and FR-TM. Moreover, the $^3$Chl *a* was observed in FDMR upon detection of the long-wavelength emission ($\lambda$>715 nm), where the emission of a Chl *a*-only PSII is only due to the vibrational tail, whereas the spectrum in that wavelength region is dominated by Chl *d*/*f* emission instead. Furthermore, a typical Chl *a*-binding PSII is expected to give rise, under the experimental conditions employed in this study, to a triplet populated by the radical pair mechanism, whereas the TR-EPR spectrum analysis indicates population by ISC. A different interpretation for the observed signal shall then be sought.

A proposition that would accommodate the just discussed $^3$Chl *a* characteristics is that of considering its formation in a subpopulation of PSII core-like complex harboring the intrinsic red-shifted Chls *d*/*f* in CP43 and CP47, but only Chl *a* in the RC (or D1/D2 complex). The ISC mechanism in this RC sub-population would then be the result of a perturbation of the cofactor energetics, for instance resulting from the specific amino acid substitution present in the far-red isoforms, that would tend to favor ISC with respect to RP recombination, similarly to the situation already discussed for the PSI reaction center of *Acaryochloris marina*[12,13]. Substantially, ISC can become the dominant mechanism if the repopulation of the RC excited state from the charge-separated radical pair kinetically outcompetes singlet-triplet mixing.

It is interesting to notice that the characteristics of the $^3$Chl *a* detected in this study, show some significant resemblance to those reported for the recombinant rChlF, where a $^3$Chl *a* is populated by the ISC mechanism and its ODMR-detected T–S spectrum also shows a maximal bleaching at ~680 nm[14]. Moreover, the ZFS and the sub-level populations of the $^3$Car detected in TR-EPR experiments are also closely comparable to those retrieved in the rChlF[14]. However, the rChlF is considered to be a homodimer of the so-called rogue psbA4 D1 isoform[32], lacking the antenna complement. Therefore, in the rChlF the $^3$Car was assigned to one of the *β*-carotene molecules that are known to be coordinated by the canonical heterodimeric D1/D2/cyt$b_{559}$ complex. The similarity of the $^3$Car properties observed in this study argues for an analogous localization. On the other hand, the overall spectral characteristic would favor a scenario in which the D1 isoform specific to ChlF is part of a modified core-like antenna harboring structure similarly to the assembly proposed by Trinugroho *et al.*[33]. Although this suggestion does certainly require further experimental support, it would also explain the substantial copurification of a core-like form of native ChlF and the FR-PSII because of their overall close structural resemblance and would, moreover, imply its constitutive presence in thylakoids and cells under conditions promoting the FaRLiP adaption response.

In conclusion, although the high sensitivity and selectivity of the ODMR technique have allowed us to resolve specific features of the $^3$Chl *a*, which is likely to play a distinct role within the FaRLiP systems, at present it remains impossible to discriminate among the proposed hypotheses regarding its origin.
